# The Hallucination Machine: A Deep-Dream VR platform for Studying the Phenomenology of Visual Hallucinations

**DOI:** 10.1101/213751

**Authors:** Keisuke Suzuki, Warrick Roseboom, David J. Schwartzman, Anil K. Seth

**Affiliations:** Sackler Centre for Consciousness Science, University of Sussex, Brighton BN1 9QJ, United Kingdom; Department of Informatics, University of Sussex, Brighton BN1 9QJ, United Kingdom

**Keywords:** Visual hallucinations, virtual reality, visual phenomenology, deep convolutional neural networks, machine learning

## Abstract

Altered states of consciousness, such as psychotic or pharmacologically-induced hallucinations, provide a unique opportunity to examine the mechanisms underlying conscious perception. However, the phenomenological properties of these states are difficult to isolate experimentally from other, more general physiological and cognitive effects of psychoactive substances or psychopathological conditions. Thus, simulating phenomenological aspects of altered states in the absence of these other more general effects provides an important experimental tool for consciousness science and psychiatry. Here we describe such a tool, the *Hallucination Machine*. It comprises a novel combination of two powerful technologies: deep convolutional neural networks (DCNNs) and panoramic videos of natural scenes, viewed immersively through a head-mounted display (panoramic VR). By doing this, we are able to simulate visual hallucinatory experiences in a biologically plausible and ecologically valid way. Two experiments illustrate potential applications of the *Hallucination Machine*. First, we show that the system induces visual phenomenology qualitatively similar to classical psychedelics. In a second experiment, we find that simulated hallucinations do not evoke the temporal distortion commonly associated with altered states. Overall, the *Hallucination Machine* offers a valuable new technique for simulating altered phenomenology without directly altering the underlying neurophysiology.

## 1.0 Introduction

There is a long history of studying altered states of consciousness (ASC) in order to better understand phenomenological properties of conscious perception ^1,2^. Altered states are defined as a *qualitative* alteration in the overall pattern of mental functioning, such that the experiencer feels their consciousness is radically different from "normal" ^1–3^, and are typically considered distinct from common global alterations of consciousness such as dreaming. ASC are not defined by any particular content of consciousness, but cover a wide range of qualitative properties including temporal distortion, disruptions of the self, ego-dissolution, visual distortions and hallucinations, among others ^4–7^. Causes of ASC include psychedelic drugs (e.g., LSD, psilocybin) as well as pathological or psychiatric conditions such as epilepsy or psychosis ^8–10^. In recent years, there has been a resurgence in research investigating altered states induced by psychedelic drugs. These studies attempt to understand the neural underpinnings that cause altered conscious experience ^11–13^ as well as investigating the potential psychotherapeutic applications of these drugs ^4,12,14^. However, psychedelic compounds have many systemic physiological effects, not all of which are likely relevant to the generation of altered perceptual phenomenology. It is difficult, using pharmacological manipulations alone, to distinguish the primary causes of altered phenomenology from the secondary effects of other more general aspects of neurophysiology and basic sensory processing. Understanding the specific nature of altered phenomenology in the psychedelic state therefore stands as an important experimental challenge.

Here, we address this challenge by combining virtual reality and machine learning to isolate and simulate one specific aspect of psychedelic phenomenology: visual hallucinations. In machine learning, deep neural networks (DNNs) developed for machine vision have now improved to a level comparable to that achieved by humans ^15,16^. For example, deep convolutional neural networks (DCNNs) have been particularly successful in the difficult task of object recognition in photographs of natural scenes ^17,18^.

Studies comparing the internal representational structure of trained DCNNs with primate and human brains performing similar object recognition tasks, have revealed surprising similarities in the representational spaces between these two distinct systems ^19–21^. For example, the neural responses induced by a visual stimulus in the human inferior temporal (IT) cortex, widely implicated in object recognition, have been shown to be similar to the activity pattern of higher (deeper) layers of the DCNN ^22,23^. Features selectively detected by lower layers of the same DCNN bear striking similarities to the low-level features processed by the early visual cortices such as V1 and V4. These findings demonstrate that even though DCNNs were not explicitly designed to model the visual system, after training for challenging object recognition tasks they show marked similarities to the functional and hierarchical structure of human visual cortices.

Trained DCNNs are highly complex, with many parameters and nodes, such that their analysis requires innovative visualisation methods. Recently, a novel visualisation algorithm called *Deep Dream* was developed for this purpose ^24,25^. *Deep Dream* works by clamping the activity of nodes at a user-defined layer in the DCNN and then inverting the information flow, so that an input image is changed until the network settles into a stable state (some additional constraints are needed, e.g. ensuring that neighbouring pixels remain strongly correlated). Intuitively, this means changing the *image* rather than changing the *network* in order to match the features of the image with what is represented in the target layer – so that the resulting image is shaped by what the network ‘expects’ to see, at the level of detail determined by the clamped layer. More precisely, the algorithm modifies natural images to reflect the categorical features learnt by the network ^24,25^, with the nature of the modification depending on which layer of the network is clamped (see Figure 1). What is striking about this process is that the resulting images often have a marked ‘hallucinatory’ quality, bearing intuitive similarities to a wide range of psychedelic visual hallucinations reported in the literature (e.g. McKenna, 2004; Shanon, 2002; Siegel & Jarvik, 1975)(see Figure 1).

**Figure 1.**
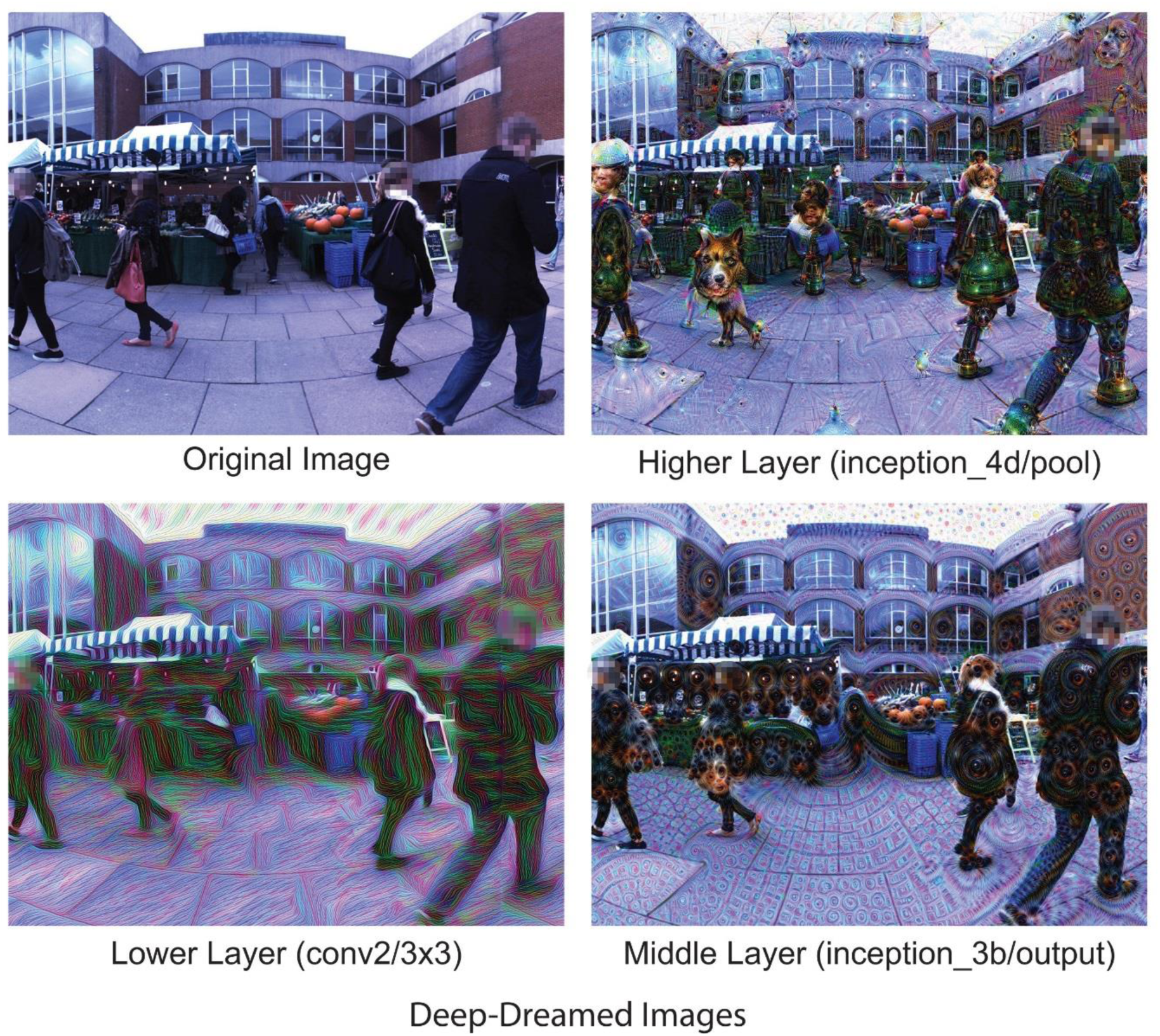
An example of the original scene (top left) and *Deep-Dreamed* scenes (top right, bottom left and right). The top right image was generated by selecting a higher DCNN layer that responds selectively to higher-level categorical features (layers = ‘inception_4d/pool’, octaves = 3, octave scale = 1.8, iterations = 32, jitter = 32, zoom = 1, step size = 1.5, blending ratio for optical flow = 0.9, blending ratio for background = 0.1, for more detail see ^48^). We used these higher-level parameters to generate the *Deep Dream* video used throughout the reported experiments. The bottom left image was generated by fixing the activity of a lower DCNN layer that responds selectively to geometric image features (layer=’conv2/3×3’, other parameters as above). The bottom right image was generated by selecting a middle DCNN layer responding selectively to parts of objects (layer=’inception_3b/output’, other parameters as above).

We set out to simulate the visual hallucinatory aspects of the psychedelic state using *Deep Dream* to produce biologically realistic visual hallucinations. To enhance the immersive experiential qualities of these hallucinations, we utilised virtual reality (VR). While previous studies have used computer-generated imagery (CGI) in VR that demonstrate some qualitative similarity to visual hallucinations ^28,29^, we aimed to generate highly naturalistic and dynamic simulated hallucinations. To do so, we presented 360-degree (panoramic) videos of pre-recorded natural scenes within a head-mounted display (HMD), which had been modified using the *Deep Dream* algorithm. The presentation of panoramic video using a HMD equipped with head-tracking (panoramic VR) allows the individual’s actions (specifically, head movements) to change the viewpoint in the video in a naturalistic manner. This congruency between visual and bodily motion allows participants to experience naturalistic simulated hallucinations in a fully immersive way, which would be impossible to achieve using a standard computer display or conventional CGI VR. We call this combination of techniques the *Hallucination Machine*.

To investigate the extent to which the *Hallucination Machine* is able to simulate natural visual hallucinations, we conducted two proof-of-concept experiments. The first experiment investigated the ecological validity of experiences produced by the *Hallucination Machine*. We compared the simulated experiences produced by the *Hallucination Machine* to unaltered control videos (see Figure 1) and to those of pharmacological psychedelic states by having participants rate their subjective experience using an ASC questionnaire developed to assess psychedelic experiences ^30,31^. In a second experiment, we investigated if the experience of the *Hallucination Machine* would also lead to a commonly reported aspect of altered states of consciousness - temporal distortion ^5,6^

## 2.0 Results

We constructed the *Hallucination Machine* by applying a modified version of the *Deep Dream* algorithm ^25^ to each frame of a pre-recorded panoramic video (Figure 1, see also Supplemental Video S1) presented using a HMD. Participants could freely explore the virtual environment by moving their head, experiencing highly immersive dynamic hallucination-like visual scenes.

### 2.1 Experiment 1: Subjective experience during simulated hallucination

In Experiment 1, we compared subjective experiences evoked by the *Hallucination Machine* with those elicited by both control videos (within subjects) and by pharmacologically induced psychedelic states ^31^ (across studies). Twelve participants took part in Experiment 1. The results are shown in Figure 2. Visual inspection of the spider chart reveals that, across all dimensions of subjective experience probed by the questionnaire, the experiences elicited by the *Hallucination Machine* are qualitatively distinct from the control videos (Fig 2a), but qualitatively similar to psilocybin experiences (Fig 2b).

**Figure 2.**
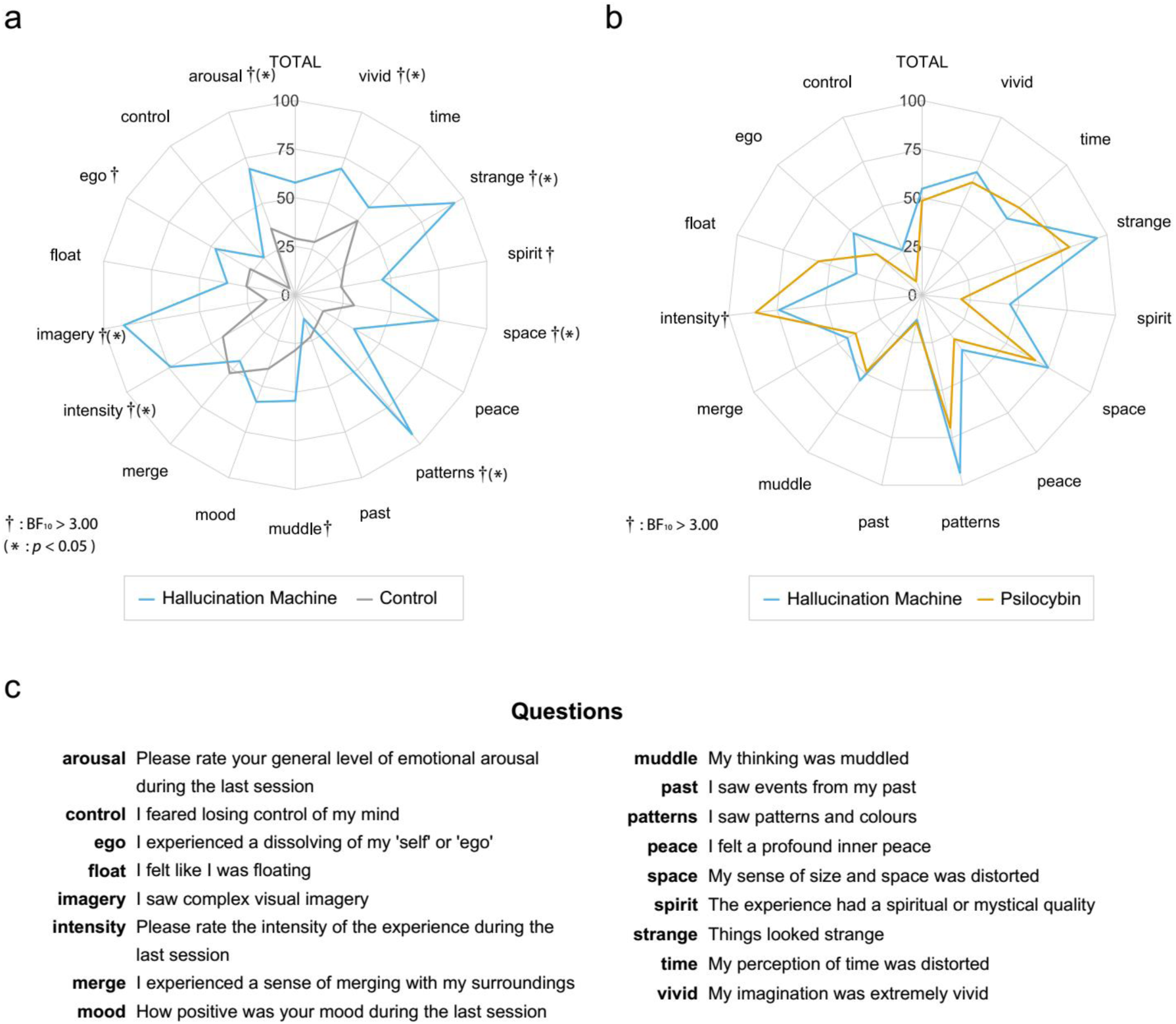
ASC questionnaire responses obtained in Experiment 1. **a**. Comparison of *Hallucination Machine* and control video responses. Stronger evidence in favour of a difference using Bayesian *t*-tests between *Hallucination Machine* and the control videos were found for ten of the questions (†: BF_10_ > 3). Standard *t*-test showed the significant differences for eight of the questions (* *p* <0.05). **b**. Comparison of *Hallucination Machine* and responses following administration of psilocybin, taken from ^31^. Bayes Factor paired sample *t*-tests revealed that responses to the question ‘intensity’ (†: BF_10_ > 3) after the *Hallucination Machine* had stronger evidence in favour of a difference from the ratings given for psilocybin experiences. **c.** Abbreviations and questions used in ASC questionnaire.

To quantify these observations, we first conducted Bayesian within-subject *t*-tests comparing responses to the ASC questionnaire following *Hallucination Machine*, and following control videos, on the null hypothesis of ‘no difference’. The analysis revealed evidence supporting the alternative hypothesis, suggesting that for the following dimensions there was a significant difference in subjective ratings between video type: ‘intensity’, ‘patterns’, ‘imagery’, ‘ego’, ‘arousal’, ‘strange’, ‘vivid’, ‘space’, ‘muddle’, ‘spirit’ (for statistics see Table 1). Bonferroni corrected, within-subject *t*-tests were consistent with the Bayesian results, with the exception of the ‘ego’, ‘muddle’, and ‘spirit’ dimensions as shown by the *p*-values in Table 1.

**Table 1.**
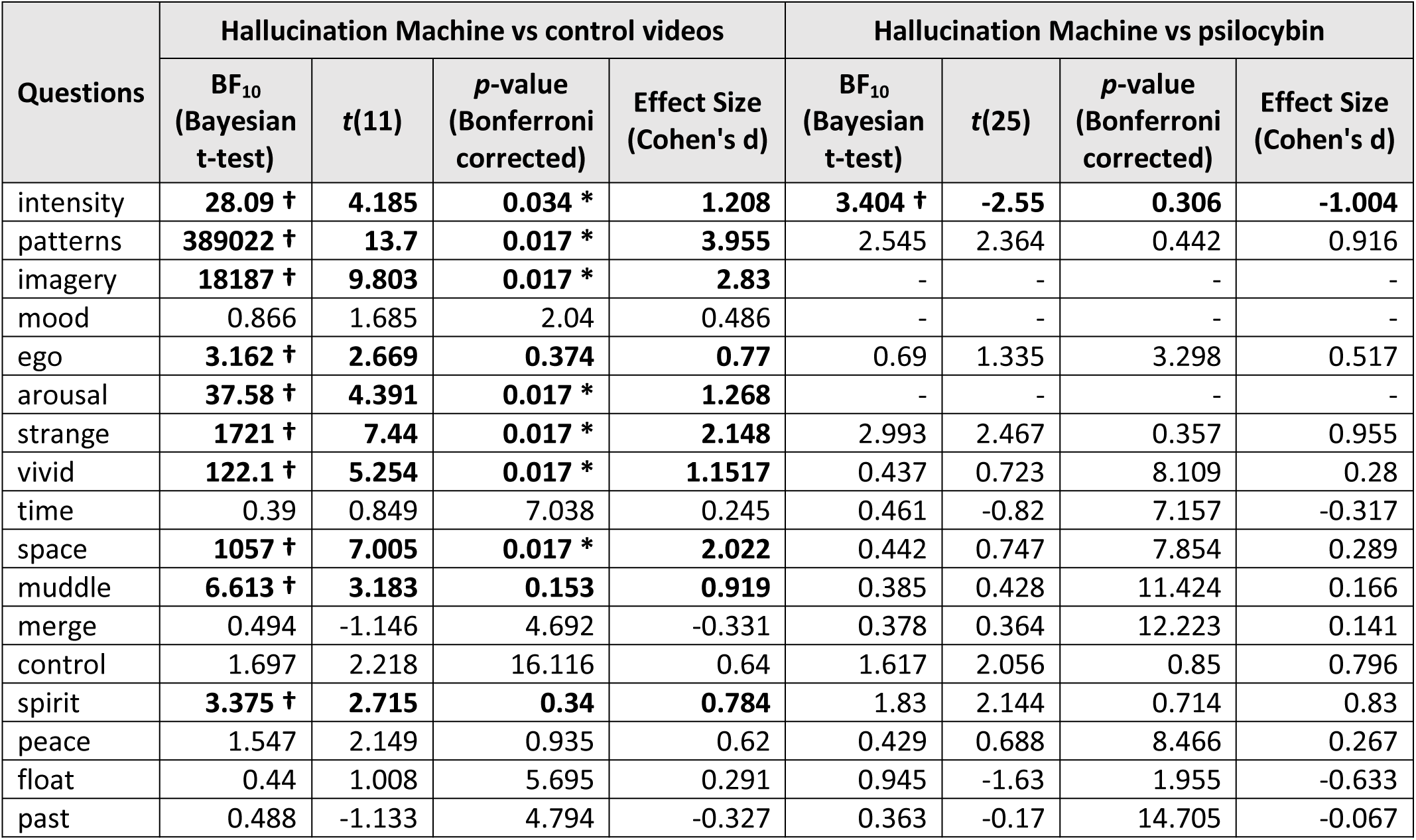
Bayesian and standard statistical comparisons of ASCQ ratings from Experiment 1 between *Hallucination Machine* and control video exposure, and between *Hallucination Machine* and psilocybin administration, data taken from ^31^. Dagger symbols and bold text indicates Bayes Factor values which show evidence in favour of a difference between ASCQ responses (†: BF_10_ > 3). Asterisks after *p*-value indicates the significant differences in standard *t*-test (* *p* <0.05). See Figure 2**c** for Abbreviations and questions used in ASCQ.

Independent Bayesian *t*-tests comparing responses to the ASC questionnaire following *the Hallucination Machine*, or following administration of psilocybin (data from a previous study^31^), also revealed evidence supporting the alternative hypothesis for the following dimension ‘intensity’, with weaker evidence for ‘pattern’ and ‘strange’, suggesting that there are some qualitative differences between *Hallucination Machine* and psilocybin experiences (see Table 1 for statistics). Crucially, for the remaining questions, Bayesian analyses were not sensitive to whether the null or alternative hypothesis was supported, but were trending in the direction of the null, i.e. no difference between subjective experiences between the *Hallucination Machine* and psilocybin: ‘vivid’ ‘time’, ‘space’, ‘muddle’, ‘peace’, and ‘past’. Standard paired *t*-test Bonferroni corrected for multiple comparisons between ASC responses following the *Hallucination Machine* and psilocybin did not reach significance for any of the question.

Together these analyses suggest that for many dimensions of subjective experience – as reflected in the ASC questionnaire - the *Hallucination Machine* induced significant changes as compared to viewing unaltered control videos, and that these changes were broadly similar to those caused by the administration of psilocybin.

### 2.2 Experiment 2: Temporal distortion during simulated hallucination

Experiment 1 showed that subjective experiences induced by the *Hallucination Machine* displayed many similarities to characteristics of the psychedelic state. Based on this finding we next used the *Hallucination Machine* to investigate another commonly reported aspect of ASC – temporal distortions^5,6^, by asking twenty-two participants to complete a temporal production task during presentation of *Hallucination Machine*, or during control videos.

One participant was excluded from the analysis due to producing intervals in the experimental session that were temporally inverted compared to the target durations. A two-way Bayesian repeated measures ANOVA consisting of factors target interval [1s, 2s, 4s] and video type (control/*Hallucination Machine*) showed the strongest evidence for an effect of target interval only (BF_10_ = 1.178 × 10^46^, 1s (*M*=1.75 s *SE*=0.09s), 2s (*M*=2.41 s *SE*=0.11 s) and 4 s (*M*=4.38 s *SE*=0.16 s)). A model including only video type showed evidence in favour of the null hypothesis (BF_10_ = 0.194), indicating that video type did not affect interval production (Figure 3). An additional two-factorial repeated measures ANOVA revealed a significant main effect of target interval (F(20,2) = 267.362, *p* < 0.01, η^2^=0.930) without the interaction (F(20, 2) = 0.935, *p*< 0.401, η^2^=0.045). However, the main effect of video type did not reach significance (F(20,1) = 0.476, *p* = 0.498, η^2^=0.023).

**Figure 3.**
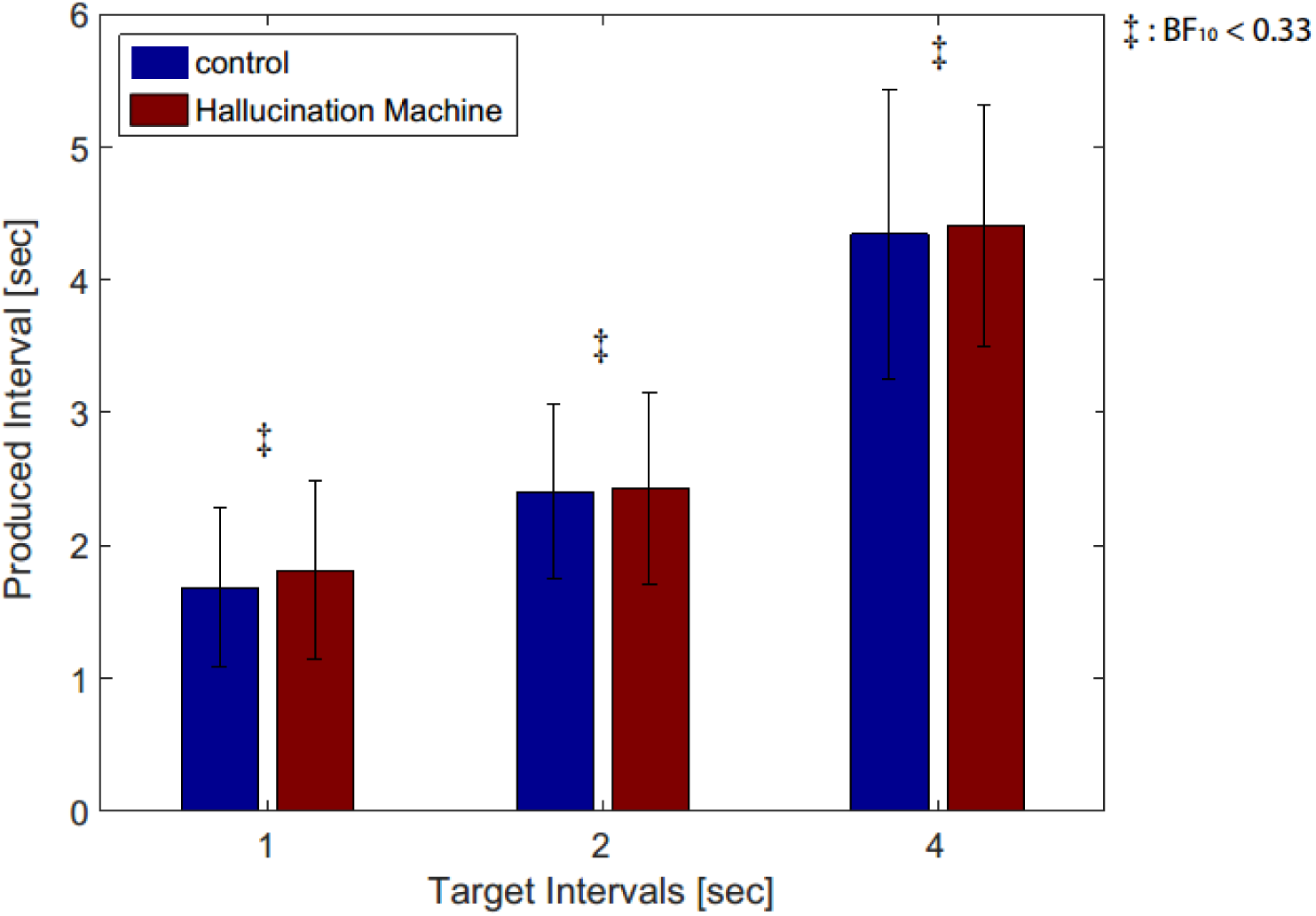
Results of temporal production task during presentation of *Hallucination Machine* or control videos. Produced time intervals are shown for both video types and target durations (1 second for low, 2 seconds for middle and 4 seconds for the high pitch tone). Bayes Factor analysis revealed strong evidence for no difference in subjective responses across video type (‡: BF_10_ < 1/3).

Post-hoc standard and Bayesian *t*-tests were applied to the participant’s subjective ratings for the six questions about their experiences during each video (see Figure 4). These revealed some differences in the *Hallucination Machine* compared to control video. Participants’ ratings of ‘presence’, “How much did you feel as if you were ‘really there’, BF_10_= 26.960, *t*(20) = 3.705, *p*=0.007, Cohen’s d=0.808; and ‘attention’, “How focused were you on the time production task”, BF_10_ = 4.830, *t*(20) = 2.822, *p* = 0.077, Cohen’s d=0.616 were reduced during the *Hallucination Machine*. Responses regarding Duration (BF_10_ = 0.278) and Speed (BF_10_ = 0.281) both revealed evidence for no difference between *Hallucination Machine* and control video. Other comparisons failed to reach an evidentiary threshold in both Bayesian and normal *t*-tests.

**Figure 4.**
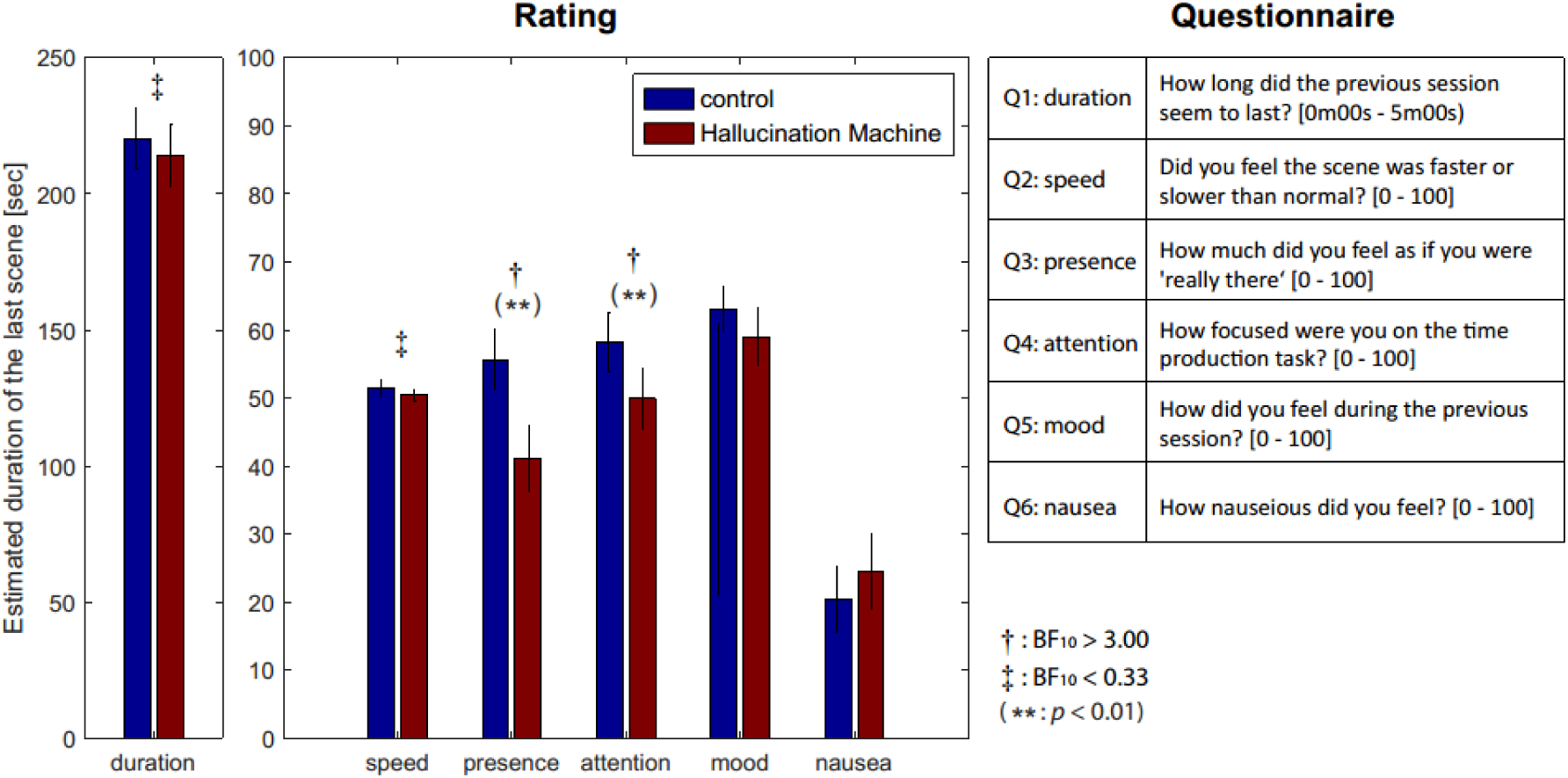
Questionnaire responses obtained in Experiment 2. Participant’s estimates’ of the total duration of *Hallucination Machine* and control videos in seconds (left panel). Participants’ subjective ratings between *Hallucination Machine* and control videos (centre). Questions used in Experiment 2 (Right). All questions were presented inside the head mounted display and participants responded to each question using a mouse to indicate their responses via a visual analog scale. Bayes Factor analysis revealed evidence in favour of a difference across video type for Q1: duration and Q2: speed (†: BF_10_ > 3), whereas evidence for no difference was found for Q3: presence and Q4: attention (‡: BF_10_ < 1/3).

## 3.0 Discussion

We have described the implementation of the *Hallucination Machine*, which provides a novel method for investigating (visual) hallucinogenic phenomenology. It combines two technologies: Panoramic video of natural scenes presented using VR, allowing the video to be experienced in a fully immersive environment, and an application of deep convolutional neural networks (DCNNs)*, Deep Dream*, which when suitably adapted can transform panoramic video to mimic hallucinatory phenomenology in a biologically plausible manner. The *Hallucination Machine* enables systematic and parameterizable manipulation of distinct aspects of altered states of consciousness (ASCs), specifically visual hallucinations, without involving the widespread systemic effects caused by pharmacological manipulations.

In two experiments we evaluated the effectiveness of this system. Experiment 1 compared subjective experiences evoked by the *Hallucination Machine* with those elicited by both (unaltered) control videos (within subjects) and by pharmacologically induced psychedelic states (across studies). Comparisons between control and *Hallucination Machine* with natural scenes revealed significant differences in perceptual and imagination dimensions (‘patterns’, ‘imagery’, ‘strange’, ‘vivid’, and ‘space’) as well as the overall intensity and emotional arousal of the experience. Notably, these specific dimensions were also reported as being increased after pharmacological administration of psilocybin ^31^.

Experiment 1 therefore showed that hallucination-like panoramic video presented within an immersive VR environment gave rise to subjective experiences that displayed marked similarities across multiple dimensions to actual psychedelic states ^31^. Although we were not able to directly compare the *Hallucination Machine* experiences to pharmacologically induced psychedelic experiences in the same subjects, the pattern of findings in Experiment 1 support the conclusion that the *Hallucination Machine* successfully simulates many aspects of ASC induced by psychedelic drugs.

Experiment 2 tested whether participants’ perceptual and subjective ratings of the passage of time were influenced during simulated hallucinations, this was motivated by subjective reports of temporal distortion during ASC ^5,6^. In contrast to these earlier findings, neither objective measures (using a temporal production task) nor subjective ratings (retrospective judgements of duration and speed, Q1 and Q2 in Figure 4) showed significant differences between the simulated hallucination and control conditions. This suggests that experiencing hallucination-like phenomenology is not sufficient to induce temporal distortions, raising the possibility that temporal distortions reported in pharmacologically induced ASC may depend on more general systemic effects of psychedelic compounds.

A crucial feature of the *Hallucination Machine* is that the *Deep Dream* algorithm used to modify the input video is highly parameterizable. Even using a single DCNN trained for a specific categorical image classification task, it is possible with *Deep Dream* to control the level of abstraction, strength, and category type of the resulting hallucinatory patterns. In the current study, we chose a relatively higher layer and arbitrary category types (i.e. a category which appeared most similar to the input image was automatically chosen) in order to maximize the chances of creating dramatic, vivid, and complex simulated hallucinations. Future extensions could ‘close the loop’ by allowing participants (perhaps those with experience of psychedelic or psychopathological hallucinations) to adjust the *Hallucination Machine* parameters in order to more closely match their previous experiences. This approach would substantially extend phenomenological analysis based on verbal report, and may potentially allow individual ASCs to be related in a highly specific manner to altered neuronal computations in perceptual hierarchies.

Another key feature of the *Hallucination Machine* is the use of highly immersive panoramic video of natural scenes presented in virtual reality (VR). Conventional CGI-based VR applications have been developed for analysis or simulation of atypical conscious states including psychosis, sensory hypersensitivity, and visual hallucinations ^28,29,32–34^. However, these previous applications all use of CGI imagery, which while sometimes impressively realistic, is always noticeably distinct from real-world visual input and is therefore suboptimal for investigations of altered visual phenomenology. Our setup, by contrast, utilises panoramic recording of real world environments thereby providing a more immersive naturalistic visual experience enabling a much closer approximation to altered states of visual phenomenology. In the present study, these advantages outweigh the drawbacks of current VR systems that utilise real world environments, notably the inability to freely move around or interact with the environment (except via head-movements).

Besides having potential for non-pharmacological simulation of hallucinogenic phenomenology, the *Hallucination Machine* may shed new light on the neural mechanisms underlying physiologically-induced hallucinogenic states. This potential rests on the close functional mappings between the architecture of DCNNs like those used here and the functional architecture of the primate visual system ^35^, as well as the equivalences between the ‘top-down’ functional flow (back propagation in *Deep Dream*) of the *Deep Dream* algorithm and the role of top-down signalling in Bayesian or ‘predictive processing’ theories of perception^36^.

A defining feature of the *Deep Dream* algorithm is the use of backpropagation to alter the input image in order to minimize categorization errors. This process bears intuitive similarities to the influence of perceptual predictions within predictive processing accounts of perception. In predictive processing theories of visual perception, perceptual content is determined by the reciprocal exchange of (top-down) perceptual predictions and (bottom-up) perceptual predictions errors. The minimisation of perceptual prediction error, across multiple hierarchical layers, approximates a process of Bayesian inference such that perceptual content corresponds to the brain’s “best guess” of the causes of its sensory input. In this framework, hallucinations can be viewed as resulting from imbalances between top-down perceptual predictions (prior expectations or ‘beliefs’) and bottom-up sensory signals. Specifically, excessively strong relative weighting of perceptual priors (perhaps through a pathological reduction of sensory input, see (Abbott, Connor, Artes, & Abadi, 2007; Yacoub & Ferrucci, 2011)) may overwhelm sensory (prediction error) signals leading to hallucinatory perceptions ^37–42^.

Close functional and more informal structural correspondences between DCNNs and the primate visual system have been previously noted ^20,35^. Broadly, the responses of ‘shallow’ layers of a DCNN correspond to the activity of early stages of visual processing, while the responses of ‘deep’ layers of DCNN correspond to the activity of later stages of visual processing. These findings support the idea that feedforward processing through a DCNN recapitulates at least part of the processing relevant to the formation of visual percepts in human brains. Critically, although the DCNN architecture (at least as used in this study) is purely feedforward, the application of the *Deep Dream* algorithm approximates, at least informally, some aspects of the top-down signalling that is central to predictive processing accounts of perception. Specifically, instead of updating network weights via backpropagation to reduce classification error (as in DCNN training), *Deep Dream* alters the input image (again via backpropagation) while clamping the activity of a pre-selected DCNN layer. The network itself is not altered in this process. Therefore, the result of the Deep Dream process can be intuitively understood as the imposition of a strong perceptual prior on incoming sensory data, establishing a functional (though not computational) parallel with the predictive processing account of perceptual hallucinations given above.

What determines the nature of this heterogeneity and shapes its expression in specific instances of hallucination? The content of the visual hallucinations in humans range from coloured shapes or patterns (simple visual hallucinations) ^7,43^, to more well-defined recognizable forms such as faces, objects, and scenes (complex visual hallucinations)^44,45^. As already mentioned, the output images of *Deep Dream* are dramatically altered depending on which layer of the network is clamped during the image-alteration process. Fixing higher layers tends to produce output similar to more complex hallucinations (Figure 5c, Higher Layer, see also Supplemental Video S1), while fixing lower layers tends create output images better resembling simpler geometric hallucinations (Figure 5b, Lower layer, see also Supplemental Video S2 and S3). These observations, together with the functional and structural correspondences between DCNNs and the primate visual hierarchy, is consistent with the idea that the content of visual hallucinations in humans may be shaped by the specificity with which a particular drug (or pathology) influences activity at different levels of processing within the visual hierarchy. Some example scenarios are schematically illustrated in Figure 5. In comparison to normal (veridical) perception (Figure 5a), simple kaleidoscopic phenomenology - which is somewhat characteristic of psychedelic states ^7,43^ - could be explained by increased influence of lower layers of the visual system during the interpretation of visual input, in the absence of contributions from higher categorical layers (Figure 5b). Conversely, complex visual hallucinations could be explained by the over emphasis of predictions from higher layers of the visual system, with a reduced influence from lower-level input (Figure 5c).

**Figure 5.**
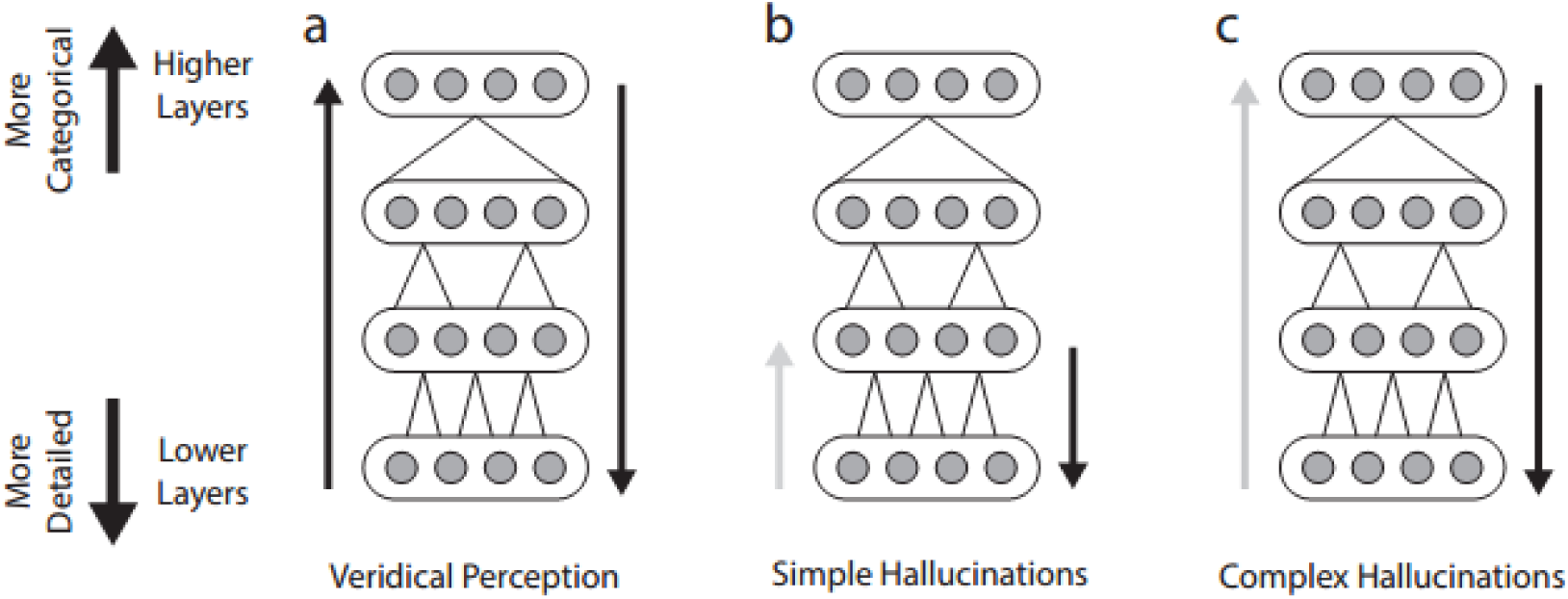
Possible hierarchical contributions to simple and complex visual hallucinations. **a.** Veridical Perception: Balanced bottom-up and top-down contributions from all levels of the hierarchy. **b**. Simple Hallucinations: perceptual content is overly influenced by visual predictions at lower network levels, with a reduced influence from lower-level input (grey arrow), emphasising features like edges and lines. **c**. Complex Hallucinations: perceptual content is overly influenced by visual predictions at higher network levels, with a reduced influence from lower-level input (grey arrow), emphasising complex object-based features.

## 4.0 Conclusion

We have described a method for simulating altered visual phenomenology similar to visual hallucinations reported in the psychedelic state. Our *Hallucination Machine* combines panoramic video and audio presented within a head-mounted display, with a modified version of ‘Deep Dream’ algorithm, which is used to visualize the activity and selectivity of layers within DCNNs trained for complex visual classification tasks. In two experiments we found that the subjective experiences induced by the *Hallucination Machine* differed significantly from control (non-‘hallucinogenic’) videos, while bearing phenomenological similarities to the psychedelic state (following administration of psilocybin). The immersive nature of our paradigm, the close correspondence in representational levels between layers of DCNNs and the primate visual hierarchy along with the informal similarities between DCNNs and biological visual systems, together suggest that the *Hallucination Machine* is capable of simulating biologically plausible and ecologically valid visual hallucinations. In addition, the method carries promise for isolating the network basis of specific altered visual phenomenological states, such as the differences between simple and complex visual hallucinations. Overall, the *Hallucination Machine* provides a powerful new tool to complement the resurgence of research into altered states of consciousness.

## 5.0 Methods

### 5.1 Hallucination Machine

In brief, the *Hallucination Machine* was created by applying the *Deep Dream* algorithm to each frame of a pre-recorded panoramic video presented using a HMD (Figure 1). Participants could freely explore the virtual environment by moving their head, experiencing highly immersive dynamic hallucination-like visual scenes.

#### 5.1.1 Panoramic video and presentation

The video footage was recorded on the University of Sussex campus using a panoramic video camera (Point Grey, Ladybug 3). The frame rate of the video was 16 fps at a resolution of 4096 × 2048. All video footage was presented using a head mounted display (Oculus Rift, Development Kit 2) using in-house software developed using Unity3D.

#### 5.1.2 DCNN specification and application of Deep Dream

The DCNN – a deeply layered feedforward neural network – used in this study had been pre-trained on a thousand categories of natural photographs used in the Large Scale Visual Recognition Challenge 2010 (ILSVRC2010) ^17,46^. During this training procedure, features in all layers are learned via backpropagation (with various modifications) to associate a set of training images to distinct categories. Consequently, the trained network implements a mapping from the pixels of the input image to the categories, represented as activation of specific units of the top layer of the network. Given this network, to create the panoramic video we applied the *Deep Dream* algorithm frame-by-frame to the raw video footage.

The *Deep Dream* algorithm also uses error backpropagation, but instead of updating the weights between nodes in the DCNN, it fixes the weights between nodes across the entire network and then iteratively updates the input image itself to minimize categorization errors via gradient descent. Over multiple iterations this process alters the input image, whatever it might be (e.g., a human face), so that it encompasses features that the layer of the DCNN has been trained to select (e.g., a dog). When applied while fixing a relatively low level of the network, the result is an image emphasizing local geometric features of the input. When applied while fixing relatively high levels of the network, the result is an image that imposes object-like features on the input, resembling a complex hallucination. Examples of the output of *Deep Dream* used in Experiments 1 and 2 are shown in Figure 1.

Although the original *Deep Dream* program was intended to process a single static image (Mordvintsev, Tyka, et al., 2015), others have developed implementations of this algorithm that process image sequences in order to make videos by blending the hallucinatory content of the previous frame with the current frame (Roelof, 2015; Samim, 2015). The principle here is to take a user defined proportion from 0–1 (blending ratio) of the previous frame’s hallucinatory patterns (0 = no information, 1 = all information) and integrate it into the current frame. In this way, each frame inherits some of the hallucinatory content of the previous frame, as opposed to *Deep Dream* starting from scratch for each frame. This frame-to-frame inheritance enables the hallucinatory patterns to remain relatively constant as the video unfolds. We extended one such implementation ^47^ to optimise the hallucinogenic properties of the video. In our extension, the optical flow of each frame is calculated by comparing the difference in the movement of all pixels between the current and previous frame. The hallucinatory patterns from areas where the optical flow was detected is merged to the current (not-yet-hallucinatory) frame based on the weighting provided by the blending ratio. The *Deep Dream* algorithm is then applied to this merged frame. We also optimised the blending ratio between each pair of frames, setting different blending ratios in areas of the image with high (foreground, moving areas, blending ratio of 0.9) or low (background static areas, blending ratio of 0.1) optical flow. This was done to avoid saturation of areas of the image with low optical flows by the higher blending ratios used for areas with high optical flow. The details of our implementation of *Deep Dream* are provided in the supplemental material. Our software for creating the *Deep Dream* video can be found on GitHub ^48^. The Deep Dream video used throughout the reported experiments was generated by selecting a higher DCNN layer, which responds selectively to higher-level categorical features (layers = ‘inception_4d/pool’, octaves = 3, octave scale = 1.8, iterations = 32, jitter = 32, zoom = 1, step size = 1.5, blending ratio for optical flow = 0.9, blending ratio for background = 0.1).

### 5.2 Experiment 1: Subjective experience during simulated hallucination

#### 5.2.1 Participants

Twelve participants completed Experiment 1 (mean age = 21.1, SD = 2.23; 7 female). Participants provided informed consent before taking part and received £10 or course credits as compensation for their time. All methods were carried out in accordance with approved guidelines provided by the University of Sussex, Research Ethics Committee.

#### 5.2.2 Experimental Design

Both experiments were performed in a dedicated VR lab. Participants were fitted with a head-mounted display before starting the experiment and exposed, in a counter-balanced manner, to either the *Hallucination Machine* or the original unaltered (control) video footage. Each video presentation lasted 3 minutes and was repeated twice, with a 180-degree direction flip of the initial orientation between the two presentations (presenting the part of the scene that would have been directly behind their viewpoint in the first presentation) to help ensure that participants experienced the majority of the panoramically-recorded scene. Participants were encouraged to freely investigate the scene in a naturalistic manner. While sitting on a stool they could explore the video footage with 3-degrees of freedom rotational movement. While the video footage is spherical, there is a bind spot of approximately 33-degrees located at the bottom of the sphere due to the field of view of the camera. After each video, participants were asked to rate their experiences for each question via an ASC questionnaire which used a visual analog scale for each question (see Figure 2c for questions used). We used a modified version of an ASC questionnaire, which was previously developed to assess the subjective effects of intravenous psilocybin in fifteen healthy human participants ^31^. All data referring to Psilocybin was taken from this study^31^.

#### 5.2.3 Analysis

Bayesian paired *t*-tests were used to compare ASC questionnaire subjective ratings between the control condition and the *Hallucination Machine*, while Bayesian independent *t*-tests were used to compare *Hallucination Machine* with subjective ratings following psilocybin administration (data taken from the original study ^31^). We quantified how close to the null (no difference between results), or to the alternative hypothesis (difference in results), each result was using JASP ^49^ with a default Cauchy prior of .707 half-width at half-maximum ^50^. A BF_10_ > 3.0 is interpreted as evidence for accepting the alternative hypothesis (i.e. there is a difference), whereas BF_10_ < 1/3 is interpreted as evidence for accepting the null hypothesis (i.e. there is no difference)^51^. Standard paired *t*-test Bonferroni corrected for multiple comparisons were also conducted.

### 5.3 Experiment 2: Temporal distortion during simulated hallucination

#### 5.3.1 Participants

A new group of Twenty-two participants that did not participate in Experiment 1 completed Experiment 2 (*M* _age_ =23.9, *SD* =6.71, 13 female). Participants provided informed consent before taking part and received £10 or course credits as compensation for their time. All methods were carried out in accordance with approved guidelines provided by the University of Sussex, Research Ethics Committee.

#### 5.3.2 Experimental Design

The experiment began with a practice session of a standard temporal production task. In each of 20 trials, participants heard one of three tones, each of a different pitch (low: 220Hz, middle: 440Hz, and high: 1760Hz, each lasting 250 milliseconds). On each trial the pitch was randomly selected. Participants were asked to produce specific time intervals for each tone (1 second for low, 2 seconds for middle and 4 seconds for the high pitch tone)^52,53^. Participants were instructed to respond immediately after the tone had ceased by holding the left mouse button down for the target time interval for each specific tone (Figure 6). After producing the interval, they were shown both their produced interval, and the target interval, as two dots on a one-dimensional scale, as well as a numeric representation (e.g. produced interval “2.4 seconds”, target interval “2.0 seconds”). The average numbers of tones per practice session was 6.12 (*SD* = 1.96) Low, 6.54 (*SD* = 1.61) Middle, and 6.33 (*SD* = 2.18) High. Participants had to repeat the practice session if the Pearson’s correlation between the target and produced intervals was less than 0.5.

**Figure 6.**
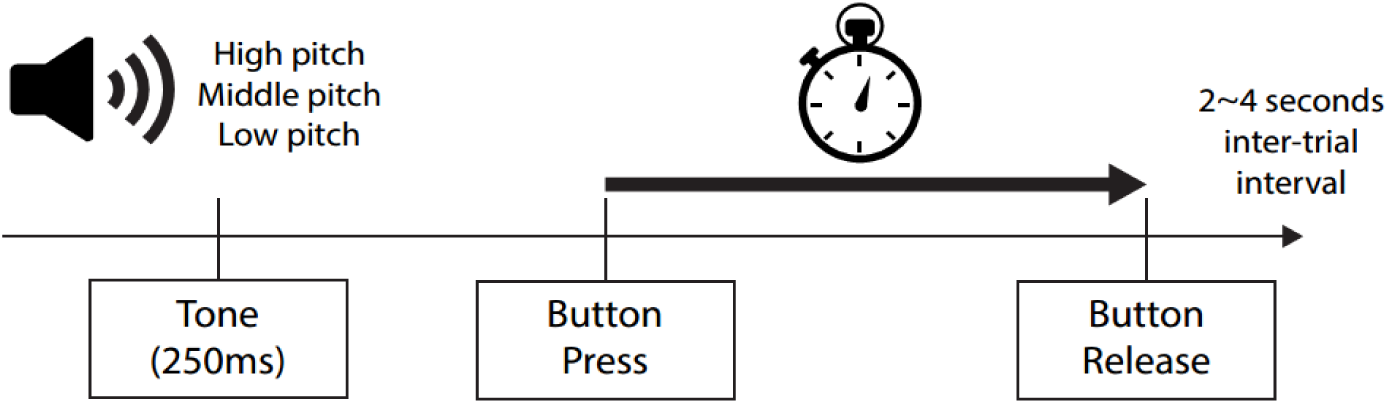
Experiment 2 temporal production task structure. While viewing either panoramic *Hallucination Machine* or control videos, participants were asked to produce one of three specific time intervals. Each time interval had been associated with a differing pitch tone during a practice session (1 second for low, 2 seconds for middle and 4 seconds for the high pitch tone). Participants responded immediately after the tone had ceased by holding the left mouse button down for the target time interval for each specific tone. After the button was released there was an inter-trial interval of between 2–4 seconds.

Once the practice was finished, participants began the experimental session. This consisted of 12 blocks. In each block a panoramic video was shown; either the control video (6 blocks) or the *Hallucination Machine* (6 blocks), and similar to Experiment 1, participants were instructed to explore the scene freely in a naturalistic manner. The order of the videos was counter-balanced across participants. Each block lasted 3 minutes, leading to a total exposure of 18 minutes for each video type. While participants explored the immersive video, low, middle or high pitch tones were presented in a random order (the average numbers of tones per block were 6.17 (*SD* = 2.02) Low, 6.16 (*SD* = 2.00) Middle, 6.21 (*SD* = 1.94) High for *Hallucination Machine*, and 6.62 (*SD* = 2.44) Low, 6.63 (*SD* = 2.42) Middle, and 6.61 (*SD* = 2.55) High for the control video). Immediately after hearing the tone, participants had to produce the interval relating to the tone (one second, two seconds, or four seconds) (Figure 6). Following the participant’s response there was a random inter-trial interval of between 2 and 4 seconds (uniformly distributed). After each block, participants answered six questions about their experiences during the video (Figure 4). The questions were presented inside the head mounted display and participants responded to the questions using a mouse to indicate a value on a visual analog scale.

#### 5.3.3 Analysis

A Bayesian two-factorial repeated measures ANOVA consisting of the factors interval production [1s, 2s, 4s] and video type (control/*Hallucination Machine*) was used to investigate the effect of video type on interval production. A standard two-factorial repeated measures ANOVA using the same factors as above was also conducted.

A two-factorial repeated measures ANOVA consisting of the factors interval production [1s, 2s, 4s] and video type (control/*Hallucination Machine*) was used to investigate the effect of video type on interval production. Similar to Experiment 1, for cases in which standard statistics did not reveal a significant difference, we quantified how close to the null (no difference between results) or alternative hypothesis (difference in results) each result was by an additional two-way Bayesian ANOVA using the same factors as above. In a similar fashion, for cases in which standard *t*-tests did not reveal significant differences in subjective ratings between video type we used additional Bayesian *t*-tests.

## Data Availability

Video materials used in the study are available in the supplemental material. The datasets generated in Experiment 1 and 2 are available from the corresponding author upon request.

## Acknowledgements

K.S., D.J.S., and A.K.S. are grateful to the Dr. Mortimer and Theresa Sackler Foundation, which supports the Sackler Centre for Consciousness Science. W.R. is supported by EU FET Proactive grant TIMESTORM: Mind and Time: Investigation of the Temporal Traits of Human-Machine Convergence. We would also like to thank Ed Venables for his help with data collection and Benjamin Ador for assistance creating the radar plot.

## Author Contributions

K.S., W.R., D.S., A.K.S., conceived and designed the study. K.S. created the materials and developed the *Hallucination Machine* system. K.S. W.R. and D.S. designed and carried out the analyses and statistical testing. K.S., D.S., and A.K.S. designed Experiment 1 and recorded the data. K.S., W.R., and A.K.S. designed Experiment 2 and recorded the data. K.S., W.R., D.S., A.K.S. wrote the manuscript together.

## Additional Information

### Competing Interests

The authors declare no competing financial interests.

